# Anti-CD38-Targeted Piperazine-Derived Lipid Nanoparticles Overcome Hepatic Clearance for mRNA Delivery to Multiple Myeloma Cells *In Vivo*

**DOI:** 10.1101/2025.11.12.687398

**Authors:** Christian G. Figueroa-Espada, Hannah C. Geisler, Ryann A. Joseph, Adele S. Ricciardi, Ann E. Metzloff, Joshua O. Acosta-González, Lulu Xue, Hannah M. Yamagata, Ajay S. Thatte, Michael J. Mitchell

## Abstract

Multiple myeloma (MM) is an incurable plasma cell malignancy characterized by clonal heterogeneity, immune evasion, and therapeutic resistance. Messenger RNA (mRNA) therapeutics offer programmable strategies to express therapeutic proteins and gene editors, but their efficacy is limited by poor extrahepatic delivery. To overcome these barriers, we developed a lipid nanoparticle (LNP) platform for targeted mRNA delivery to MM cells *in vivo*. Through combinatorial screening, we identified C16-O1, a piperazine-based ionizable lipid that efficiently transfects both CD138+ and therapy-resistant CD138-MM subclones. For tumor selectivity, LNPs were functionalized with an antibody fragment against CD38, a clinically validated MM antigen. Anti-CD38 LNPs reached the tumor-site and significantly reduced hepatic accumulation in murine xenografts. As an *in vitro* proof-of-concept, delivery of Cas9 mRNA and an IRF4-targeting guide RNA induced gene knockout, cell-cycle arrest, and lenalidomide sensitization. Together, these findings establish a robust framework for targeted mRNA delivery in MM and other hematologic malignancies.

## INTRODUCTION

Multiple myeloma (MM) is an incurable hematologic malignancy characterized by the clonal expansion of malignant plasma cells within the bone marrow.^1^ Despite significant advances in systemic therapies, patients face persistent disease relapse and shorter survival rates.^2^ This therapeutic failure is often driven by antigen escape and the emergence of phenotypically diverse, resistant subclones, including CD138^-^ cells with stem-like features and therapeutic resistance.^3–5^ These biological challenges are compounded by the unique architecture and immunosuppressive microenvironment of the bone marrow. Accordingly, there is a pressing need for flexible therapeutic platforms capable of targeting diverse myeloma subpopulations to overcome resistance mechanisms and enable synergistic combination treatments.

Messenger RNA (mRNA) therapeutics offer highly adaptable tools for transiently expressing diverse payloads, including gene editors, immune effectors, or synthetic proteins that can be used to re-sensitize tumor cells or enhance the efficacy of immunotherapies.^6–8^ However, the successful clinical application of this technology faces significant intrinsic hurdles: mRNA is an unstable molecule that is rapidly degraded by extracellular nucleases, and its large, highly anionic structure prevents efficient passage across the non-polar cellular membrane.^9,10^ These biological and physical challenges require a robust delivery system to protect the cargo, enable cellular uptake, and facilitate endosomal escape. The breakthrough success of lipid nanoparticles (LNPs) in delivering mRNA—most notably demonstrated by the rapid development of COVID-19 vaccines—has thus established them as the gold standard delivery vehicle for this versatile therapeutic platform.^11,12^

However, their use in oncology is significantly constrained by delivery challenges. Conventional LNPs are hindered by intrinsic liver tropism and a lack of active targeting, severely restricting their ability to deliver functional mRNA selectively to solid or hematologic tumors outside the liver.^13–17^ Overcoming this dominant hepatic clearance and achieving selective, functional mRNA delivery to the tumor site remains a critical unmet need in nanomedicine.

Here, we developed a programmable LNP platform designed for antigen-directed mRNA delivery to MM. The platform is based on a novel ionizable lipid, C16-O1, which was identified through a screening of chemically distinct formulations to ensure optimal delivery efficiency across the heterogeneous MM populations, specifically targeting both the bulk CD138+ cells and the resistant CD138-subclones. The modularity of this platform is key for overcoming common resistance mechanisms, such as antigen loss, by allowing for rapid exchange of targeting ligands or therapeutic payloads. We achieved selective *in vivo* targeting by conjugating the LNP surface with Fab’ fragments of an anti-CD38 monoclonal antibody via thiol-maleimide chemistry.^18^ CD38 is a clinically validated myeloma antigen that remains broadly expressed even in relapsed tumors.^19^ In murine xenograft models, these CD38-targeted C16-O1 LNPs demonstrated enhanced tumor accumulation and a significant reduction in off-target hepatic sequestration relative to untargeted controls. As an *in vitro* proof-of-concept for the flexible cargo capacity, we further demonstrated that the platform can effectively deliver Cas9 mRNA and an sgRNA targeting the transcription factor IRF4, which resulted in effective gene knockout that sensitized myeloma cells to lenalidomide.^20^ These findings establish a versatile, antigen-directed LNP platform that integrates a screened ionizable lipid and surface functionalization, enabling programmable mRNA delivery to overcome anatomical and cellular barriers in hematologic malignancies.

### Design and Physicochemical Characterization of a Piperazine-Derived Ionizable LNP Library

To engineer LNPs for mRNA delivery to MM cells *in vitro*, we formulated a library of LNPs, each with its unique ionizable lipid. Specifically, a library of ionizable lipids was synthesized, as previously described,^10^ using a fast and straightforward S_N_2 reaction setup, where one of three epoxide tails – C12, C14, or C16 – was reacted with one of three polyamine cores – O1, O2, or O3 (**Figure 1a**). Liquid chromatography-mass spectrometry (LC-MS) was used to confirm the molecular identity of these lipids, including C16-O1 (**Figure 1b, S1**).

**Figure 1.**
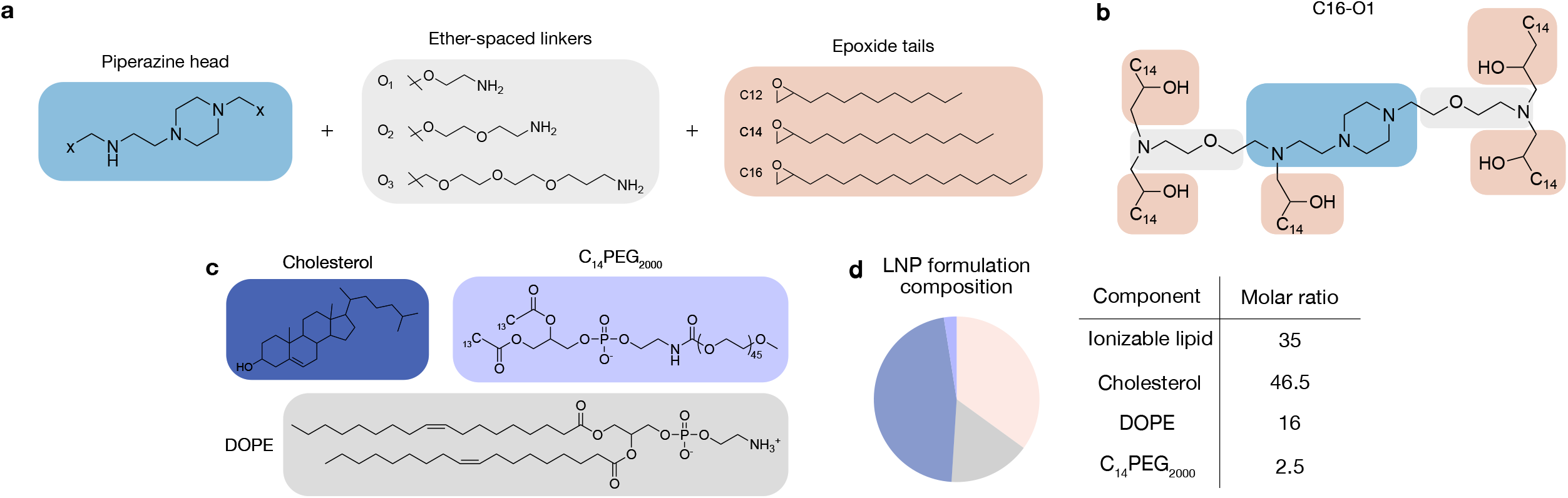
Overview of the piperazine-derived ionizable lipids and LNP library. (a) Structures of the piperazine head, the corresponding ether-spaced linkers (O1, O2, O3), and the three epoxide tails (C12, C14, C16) that were used as building blocks. These were reacted to produce nine unique ionizable lipids (C_N_-O_M_). (b) Chemical structure of piperazine-derived ionizable lipid, C16-O1, synthesized via an S_N_2 reaction. (c) Chemical structures of the other excipients that comprise the LNP formulation: cholesterol, DOPE, and PEG-lipid (C14PEG2000). (d) Pie chart and table detailing the molar ratio of each component in the standard LNP formulation used for screening every ionizable lipid in this study.

Subsequently, these lipids were individually combined with DOPE, cholesterol, and lipid-anchored poly(ethylene glycol) (PEG) to formulate LNPs (**Figure 1c, d**) via chaotic mixing with an aqueous phase of mCherry mRNA in a microfluidic device (**Figure 2a**).^21^ Each one of these lipid excipients plays a crucial role in LNP formulation, intracellular uptake, and delivery. The ionizable lipid enables mRNA encapsulation and endosomal escape for potent intracellular delivery; the phospholipid DOPE promotes LNP membrane formation, cholesterol enhances membrane stability, and the lipid-PEG limits rapid clearance and immune cell opsonization (**Figure 1c**).^22^ In addition to our library of nine LNPs, each with its unique ionizable lipid, we formulated two control LNPs with ionizable lipids C12-200 and SM-102, which serve as industry-standard and U.S. Food and Drug Administration (FDA)-approved ionizable lipids for comparison, respectively (**Figure S2**).

**Figure 2.**
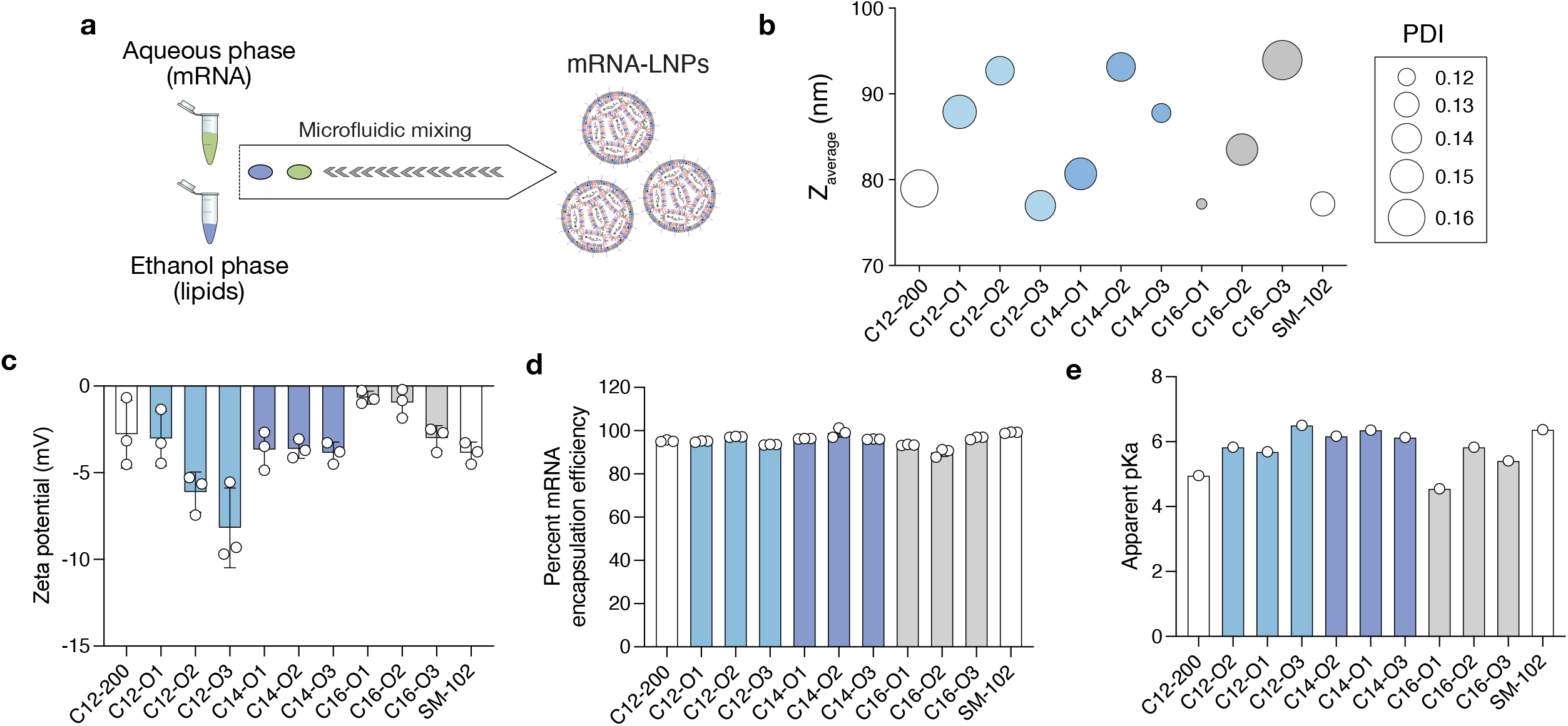
Synthesis and physicochemical characterization of ionizable LNP library. (a) Synthesis of LNPs via microfluidic mixing of an ethanol phase containing ionizable lipid, phospholipid, cholesterol, and lipid-PEG, and an aqueous phase containing mRNA. (b) Hydrodynamic diameter and polydispersity index, (c) zeta potential, (d) mRNA encapsulation efficiency, and (e) apparent pKa characterization of each LNP in the library. Data are reported as mean ± s.d.

Following formulation, we characterized the hydrodynamic size, polydispersity index (PDI), zeta potential, encapsulation efficiency, and pKa of the LNP library (**Figure 2b, c, d, e**). All nine LNPs were less than 100 nm in diameter, and all LNPs also had PDIs less than 0.17. LNPs C16-O3, C14-O2, and C12-O2 were the largest LNPs, however this did not affect mRNA encapsulation, for which all LNPs had greater than 90%. Finally, we characterized LNP pKa, or the pH at which the LNP is 50% protonated. The LNP pKa depends mainly on the ionizable lipid component. A value <7.0 is critical, as it ensures the ionizable lipid remains neutral at physiological pH (pH 7.4) but becomes protonated in the acidic environment of the endosome (pH 5.0-6.5). This protonation induces a conformational change that promotes the fusion of the LNP membrane with the endosomal membrane, thereby facilitating the release of the mRNA cargo into the cytosol. pKa values for the LNP library ranged from 4.5 to 6.5, indicating the ionizable nature of our LNPs for potent intracellular mRNA delivery.

### *In Vitro* Screening Identifies C16-O1 as an Ionizable Lipid for mRNA Delivery to Heterogeneous Myeloma Subpopulations

Next, we sought to evaluate the *in vitro* mRNA transfection efficiency of our LNP library in human MM cells. We chose three cell lines with different levels of heterogeneity in CD138 expression – NCI-H929, RPMI8226, and U266 (**Figure 3a, b**). While it is known that cell lines have differences in mimicking human disease, this number provides sufficient models to evaluate LNP-mediated mRNA delivery to MM cells *in vitro*. MM cells are characterized by having a complex genomic makeup, imposing a challenge to target them directly or enabling their ability to evade anti-cancer mechanisms for cell death.^23,24^ Notably, the CD138-population of MM cells has been linked to a high proliferative ability and a stem cell-like behavior, inducing poor response to anti-myeloma therapies.^4^ To ensure therapeutic platforms can overcome this critical hurdle, we screened our piperazine-derived ionizable LNPs for *in vitro* mRNA delivery efficiency across both CD138+ and CD138-subpopulations, with a view toward their eventual application in protein replacement or gene editing therapy.

**Figure 3.**
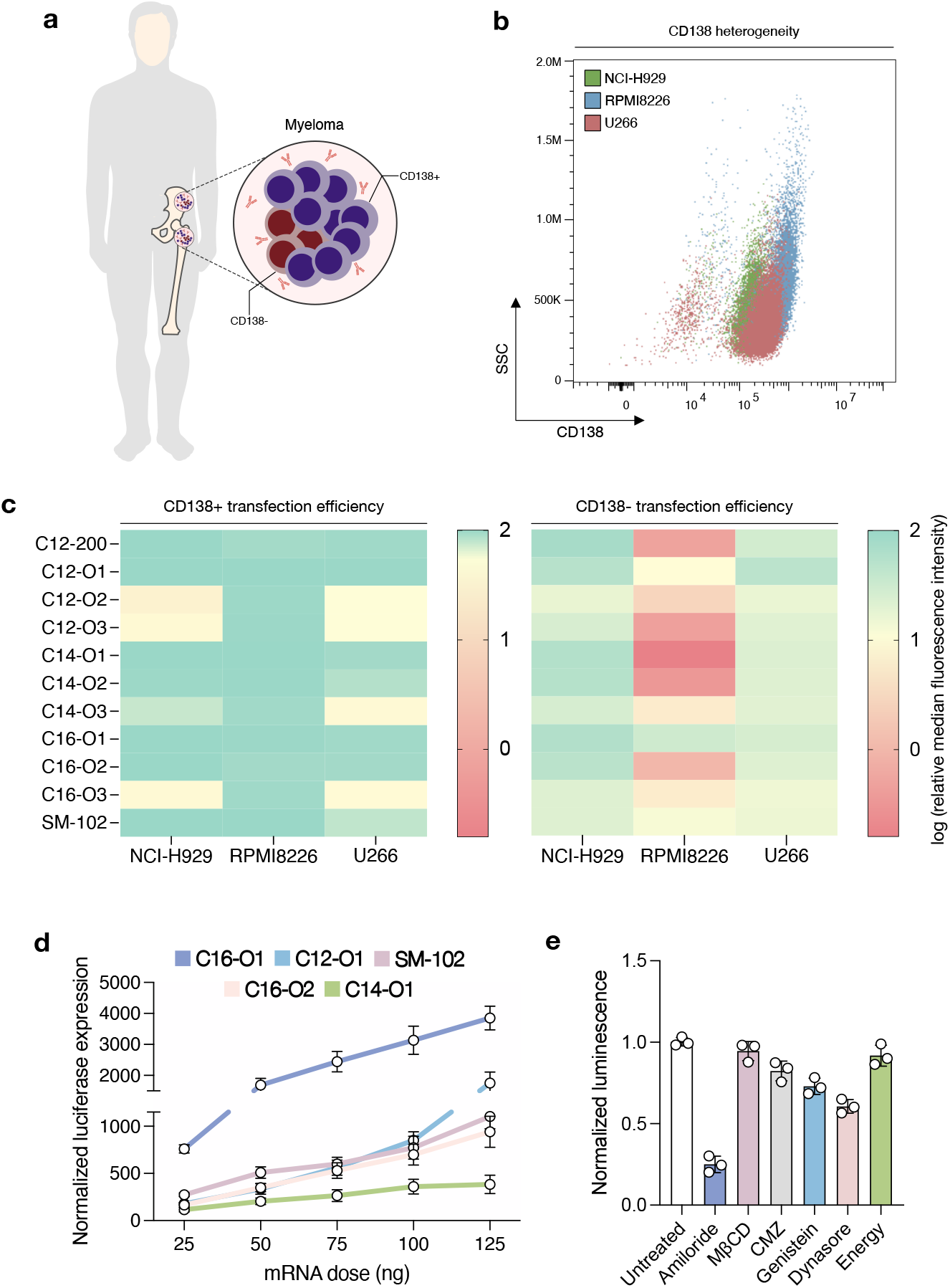
Identification of C16-O1 ionizable lipid for efficient mRNA delivery to heterogeneous MM cell subpopulations. (a) Schematic representation illustrating the diverse populations of malignant plasma cells within the bone marrow, specifically highlighting the bulk CD138+ cells and the therapy-resistant CD138-subclones. (b) Dot plot displaying the CD138 expression and side scatter (SSC) of the three human MM cell lines used in the screening (NCI-H929, RPMI-8226, and U266), confirming the heterogeneity of CD138 expression across the panel. (c) Heat maps illustrating the mCherry mRNA transfection efficiency (log relative median fluorescence intensity) of the piperazine-derived LNP library and two industry and FDA-approved standards (C12-200 and SM-102, respectively). Results are segmented by CD138+ (left) and CD138-(right). C16-O1 is identified as the lead lipid with superior efficacy across all cell lines and both subpopulations. (d) Comparison of luciferase mRNA delivery of the top-performing C16-O1 against other strong candidates (C12-O1, C16-O2, C14-O1) and the SM-102 control, demonstrating the highest dose-dependent luciferase expression for C16-O1 across a 25-ng to 125-ng range. (e) Inhibition assay demonstrating the primary internalization pathway of the C16-O1 LNP. Cells were treated with inhibitors targeting specific endocytosis routes (amiloride: macropinocytosis; MβCD: caveolae-mediated; genistein: caveolae/clathrin-mediated; dynasore: dynamin-dependent), confirming that C16-O1 uptake is primarily mediated by macropinocytosis (significantly reduced luminescence signal with amiloride treatment). Data is plotted as mean ± s.d. n = 3 biological replicates.

LNPs or Lipofectamine MessengerMAX were used to treat NCI-H929, RPMI8226, or U266 cells with 50 ng of mCherry mRNA per 60,000 cells. Lipofectamine is often considered a gold standard transfection reagent for *in vitro* nucleic acid delivery. Mean fluorescence intensity as a measure of functional mRNA delivery was evaluated in the three cell lines 24 hrs after treatment with LNPs or Lipofectamine. All eleven LNPs from the library had significantly higher mCherrry expression than Lipofectamine, which was used to normalize the data (**Figure S3a**). Of these, C16-O1 showed high efficiency for mRNA delivery across all three cell lines (**Figure 3c, d**). We also evaluated cell viability 24 hrs following LNP treatment, and none of the LNPs had a significant decrease in cell viability compared to Lipofectamine (**Figure S3b**). Lastly, we evaluated the efficiency of these LNPs to transfect both CD138+ and CD138-subpopulations of myeloma cells and found that most LNPs delivered to both populations in the NCI-H929 and U266 cell lines. However, that was not the case for RPMI-8226, where the only noticeable ionizable lipid delivering to both subpopulations was C16-O1 (**Figure S3c**). To further confirm the performance profile, we performed a dose-response study in the RPMI-8226 cells, demonstrating that C16-O1 maintained superior delivery efficiency across a range of mRNA concentrations compared to other top-performing LNPs (**Figure 3d**). Furthermore, to gain mechanistic insight into its superior function, we investigated the primary endocytic uptake route. We used pharmacological inhibitors specific to various endocytic pathways, such as clathrin- and caveolae-mediated endocytosis, and measured the subsequent impact on luciferase protein expression to define the primary route. This analysis confirmed that C16-O1 preferentially utilized the macropinocytosis pathway for cellular entry (**Figure 3e**). This dependence on macropinocytosis is a potentially advantageous feature, as this pathway is often upregulated in highly proliferative cancer cells due to oncogenic signaling.^25^ This robust performance, including superior efficiency, low cytotoxicity, a favorable micropinocytosis-dependent uptake route, and, critically, the ability to transfect the therapy-resistant CD138? subpopulation across heterogeneous cell line models, established C16-O1 as the lead ionizable LNP candidate for all subsequent studies.

### C16-O1 LNPs Deliver Gene Editing Cargo and Sensitize Myeloma Cells to Lenalidomide *In Vitro*

Proven efficient mRNA delivery to all three human myeloma cell lines, we sought to encapsulate Cas9 mRNA with an IRF4 sgRNA to evaluate whether the C16-O1 LNP platform could also be used for mRNA-mediated Cas9 genome editing. IRF4 is a transcription factor linked to myeloma cell survival, and its expression has been shown to lead to poor prognosis and unfortunate response to anti-MM therapies.^26,27^ To evaluate the potential of knocking out IRF4 with the C16-O1 LNP platform, we performed a cell cycle analysis 36 hrs after treatment with Cas9 mRNA and IRF4 sgRNA LNPs.^28^ We observed that as the total RNA dose increased, the cells were sequestered in the G2/M phase before and during mitosis (**Figure S4a**), suggesting that IRF4 knockout disrupts normal cell-cycle progression. This G2/M arrest is a known precursor to mitotic catastrophe and subsequent apoptosis in cancer cells.^29^ To mitigate concerns that the observed cytotoxicity was due to the LNP system itself, a scrambled sgRNA sequence was used as a negative control at the highest RNA dose. The lack of cell cycle disruption in this control proves that the cell death is sequence-specific and IRF4-driven, rather than an intrinsic toxicity of the C16-O1 LNP platform (**Figure S4a**).

After confirmation of IRF4 knockout following LNP treatment (**Figure S4b, c**), we sought to evaluate whether it affected how myeloma cells responded to MM chemotherapeutics, specifically lenalidomide, an FDA-approved immunomodulatory drug. Lenalidomide has pharmacological limitations, such as low solubility and bioavailability, leading to severe side effects such as rashes and gastrointestinal issues.^30^ Thus, keeping dosages of lenalidomide as low as possible while still managing the disease would be crucial for MM patients. In this study, we treated myeloma cells with the C16-O1 LNPs encapsulating Cas9 mRNA and scrambled or IRF4 sgRNA with or without lenalidomide, and evaluated cytotoxicity 7 days post-treatment. We confirmed what we had observed in the cell cycle phase analysis; an arrest before or during the mitosis phase induces cancer cell apoptosis. We observed that keeping the lenalidomide dose constant while increasing the RNA dose with the C16-O1 LNPs enhanced the killing of the cancer cells, sensitizing these to the immunomodulatory drug (**Figure S4d**). This synergistic effect confirms the dual benefit of the C16-O1 platform as it provides efficient delivery for mechanistic gene editing, which, in turn, offers a potent strategy for pharmacologic resensitization, potentially enabling the use of lower, less toxic lenalidomide doses for patients with MM.

### Targeted C16-O1 LNPs Achieve Enhanced mRNA Delivery to Myeloma Subcutaneous Tumors *In Vivo*

Following the screening of different piperazine-derived ionizable lipids and choosing to focus on C16-O1, we generated targeted LNPs. Incorporating a targeting moiety into LNPs can significantly enhance the efficiency and specificity of the delivery to hard-to-transfect cells, such as lymphocytes.^31^ To reach MM cells *in vivo* with our LNPs, we chose CD38, a glycoprotein overly expressed in MM cells and many other B-cell lymphoma cells, such as mantle cell lymphoma. CD38 is also clinically relevant for MM, specifically with Daratumumab, the first monoclonal antibody approved for treating MM.^19^ To generate the t-C16-O1, the C16-O1 LNP base formulation was modified to incorporate PEG-maleimide as one part of the molar composition of PEG (**Figure 4a**). These PEG-maleimide functionalized LNPs were then conjugated to Fab’ fragments of anti-human CD38 antibody using maleimide-thiol chemistry (**Figure 4b**). Resultant t-C16-O1 LNPs were characterized for their size, uniformity, zeta potential, and encapsulation efficiency. While changes were not observed in their uniformity, zeta potential, and mRNA encapsulation efficiency, dynamic light scattering measurements confirmed conjugation to anti-CD38 Fab’ fragments with a ~60 nm increase in the hydrodynamic diameter (**Figure 4c**).

**Figure 4.**
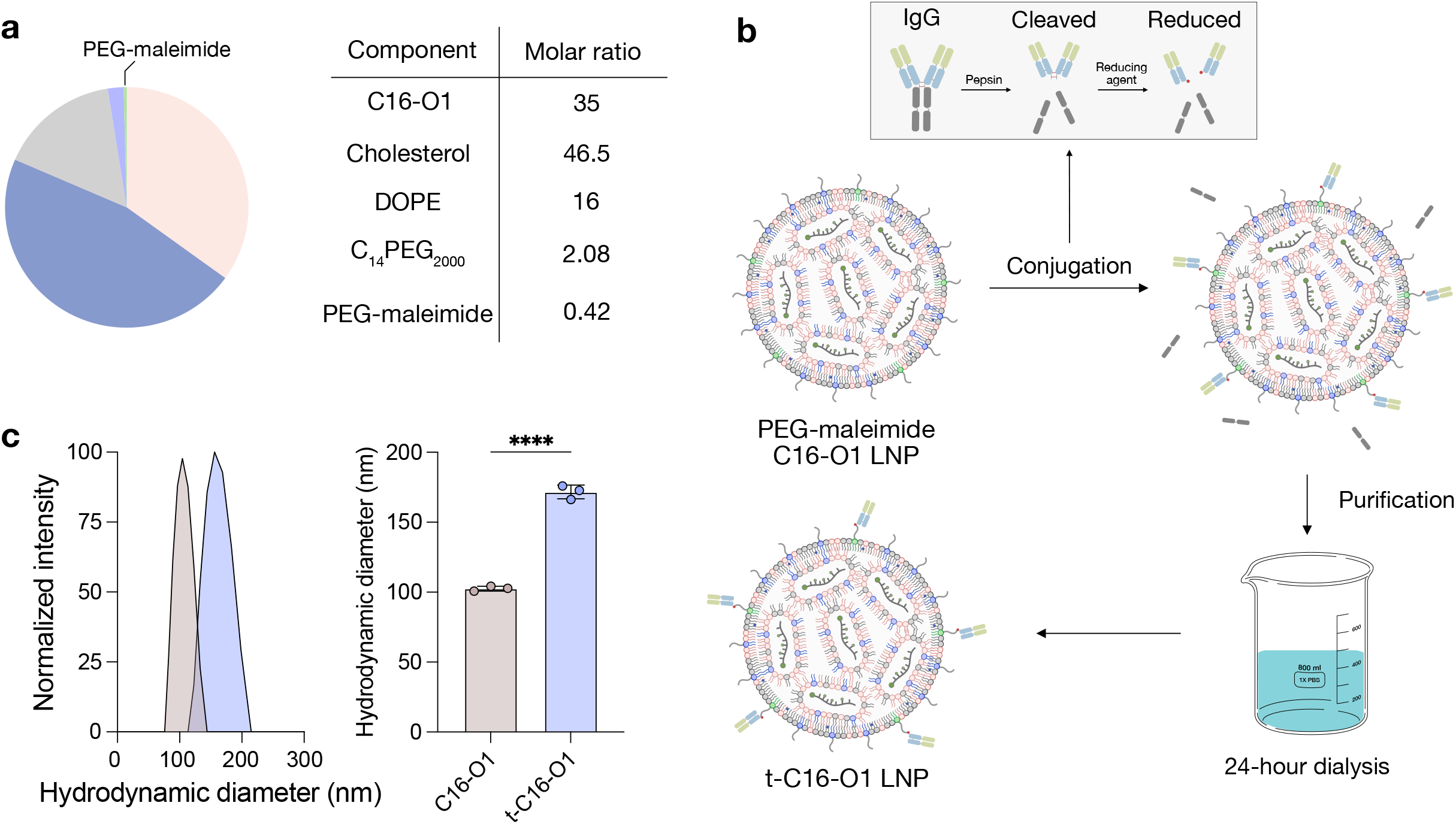
Synthesis and confirmation of t-C16-O1 mRNA LNPs. (a) Pie chart and corresponding table detailing the molar percentage of the C16-O1 LNP base formulation, including the novel C16-O1 ionizable lipid, cholesterol, DOPE, C14PEG2000 (PEG), and PEG-maleimide. The PEG-maleimide component constitutes a molar ratio of 0.42 out of the total PEG molar composition (2.5). (b) The targeting moiety is generated by cleaving the anti-human CD38 whole IgG antibody using Pepsin, followed by reduction to yield Fab’ fragments with reactive thiol groups. These Fab’ fragments are conjugated to the PEG-maleimide functionalized C16-O1 LNP surface via maleimide-thiol chemistry (thiol-maleimide conjugation). The resulting t-C16-O1 LNPs are purified via 24-hour dialysis in PBS to remove unconjugated components. (c) DLS measurement confirming successful Fab’ conjugation onto the LNP surface. The graph shows the normalized intensity distribution, and the bar plot quantifies the significant increase (~60 nm) in the hydrodynamic diameter of t-C16-O1 compared to the untargeted C16-O1 LNP. Data is plotted as mean ± s.d.

Following the generation and confirmation of t-C16-O1, we moved to *in vivo* to probe their efficacy in targeting myeloma cells in tumor-bearing mice. Although myeloma originates in the bone marrow, we utilized a subcutaneous model as it provides proof-of-concept to assess the *in vivo* targeting abilities of the t-C16-LNPs. To create the model, GFP-expressing RPMI-8226 tumor cells were inoculated in the right flank of immunocompromised mice for 21 days prior to LNP administration (Figure 5a). The biodistribution of the t-C16-O1 LNPs was compared against two controls: untargeted C16-O1 LNPs, and the FDA-approved SM-102, which showed higher delivery to CD138-MM cells than industry-standard C12-200 (**Figure S3c**). To better understand the tropism of these LNPs, we labeled them with DiR on their surface, which allowed us to image organs to detect the presence of the LNPs. As such, our experimental groups included free DiR, SM-102, C16-O1, and t-C16-O1 LNPs. The tumors, livers, lungs, and spleens were harvested 6 hours post-injection and analyzed by IVIS live imaging (**Figure 5b, c**). We observed that t-C16-O1 LNPs had 2.3-, and 3.7-fold higher tumor delivery than SM-102 and untargeted C16-O1 LNPs (**Figure 5b**), respectively.

**Figure 5.**
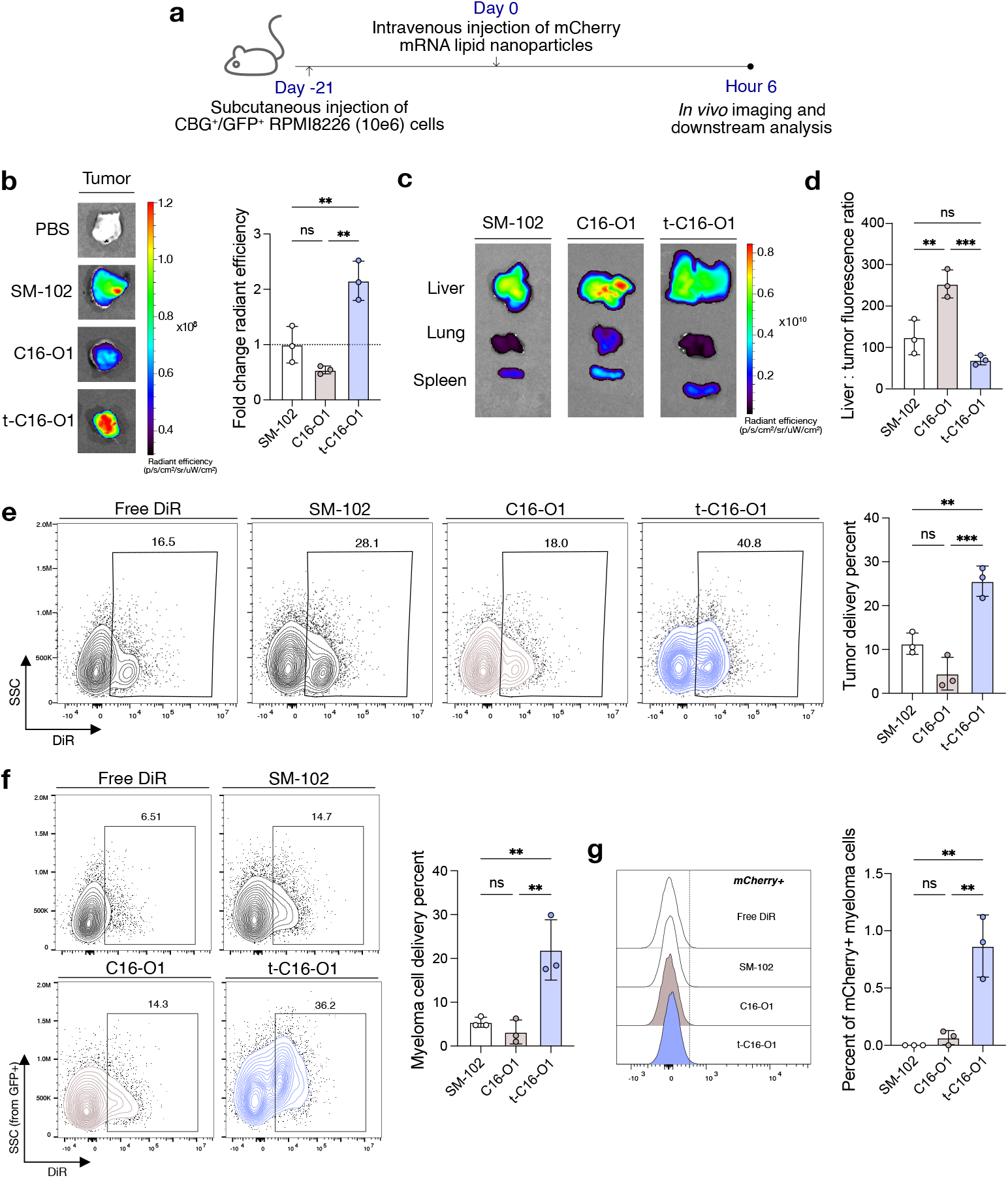
t-C16-O1 LNPs enable functional mRNA delivery to myeloma subcutaneous tumors *in vivo*. (a) Schematic of the subcutaneous RPMI-8226 xenograft model, showing tumor inoculation (GFP+ cells), LNP administration, and timeline for ex vivo analysis. (b) Representative IVIS images and corresponding radiant efficiency quantification of DiR fluorescence (LNP delivery) in tumors harvested 6 hours post-injection for SM-102, untargeted C16-O1, and targeted t-C16-O1 LNPs compared to control (free-DiR). (c) Representative IVIS images of major organs (liver, lung, spleen) and their corresponding radiant efficiency. (d) Quantification of the liver-to-tumor DiR fluorescence ratio, demonstrating reduced hepatic tropism for the t-C16-O1 LNP. (e) Contour diagrams for the different groups showing the gating strategy used to determine tumor mRNA-LNP delivery, specifically identifying DiR+ cells within the GFP+ myeloma population. (f) Myeloma cell-specific LNP uptake, quantified as the percentage of DiR+ cells within the GFP+ myeloma population. (g) Functional mRNA translation *in vivo*, quantified as the percentage of mCherry+ cells within the GFP+ myeloma population. The signal from the free-DiR groups was subtracted from the LNP treatment groups. Data is plotted as mean ± s.d. n = 3 mice per group.

We next addressed the critical challenge of hepatic clearance. While the untargeted C16-O1 LNPs demonstrated significant liver tropism, the incorporation of the CD38 targeting moiety resulted in a dramatic reduction of the liver-to-tumor delivery ratio by at least 3-fold (**Figure 5c, d**). This reduction directly demonstrates that surface functionalization successfully re-routes LNP biodistribution away from the liver, achieving enhanced and tumor-specific targeted delivery. Importantly, we observed no significant changes in serum aspartate or alanine aminotransferase (AST or ALT) levels compared to the saline control, indicating that the t-C16-O1 LNPs exhibited excellent *in vivo* safety and low systemic toxicity (**Figure S5a, b**).

To confirm efficient delivery and cellular specificity, we also evaluated LNP uptake and mRNA function in the tumor and myeloma cells using flow cytometry. We observed 26% of tumor delivery with the t-C16-O1 LNPs, while SM-102 and untargeted C16-O1 LNPs yielded 10% and 4%, respectively (**Figure 5e**). Further, we assessed myeloma cell-specific delivery gating on GFP+ cells and found both SM-102 and untargeted C16-O1 LNPs with less than 5% delivery, while t-C16-O1 LNPs have over 20% delivery (**Figure 5f**). Crucially, functional delivery, assessed by mCherry mRNA translation within GFP+ myeloma cells, confirmed the specificity of the t-C16-O1 platform. The t-C16-O1 LNPs yielded a significant population of mCherry+ myeloma cells, 1%, whereas untargeted C16-O1 and SM-102 resulted in negligible expression (**Figure 5g**).

Altogether, these *in vivo* biodistribution and cellular uptake data establish the t-C16-O1 platform as the first targeted mRNA LNP delivery system, to our knowledge, designed to overcome hepatic clearance and achieve specific, functional delivery to myeloma cells *in vivo*. Beyond addressing a critical unmet need in nanomedicine, the ability to achieve functional payload delivery to 1% of myeloma cells in the tumor provides a robust therapeutic foundation for combination strategies, such as using mRNA to transiently express tumor-sensitizing or immunomodulatory factors in concert with existing chimeric antigen receptor T-cell therapies,^32^ thereby offering a powerful strategy to circumvent antigen escape and relapse in MM.

## SUMMARY

In this study, we sought to interrogate a novel library of piperazine-derived ionizable lipids for their ability to deliver mRNA to heterogeneous CD138+ and CD138-myeloma cell subpopulations. The LNP library was screened across three human myeloma cell lines—NCI-H929, RPMI8226, and U266—and evaluated for efficient and non-toxic mCherry mRNA delivery. Seeking to identify a top-performing ionizable lipid, we employed a weighted logarithmic scale considering overall mRNA delivery efficiency across all cell lines and, critically, the efficiency in transfecting the therapy-resistant CD138 subpopulation. This screening process yielded lipid C16-O1 as the lead candidate for mRNA delivery to myeloma cells.

We next tested whether the C16-O1 LNP could efficiently encapsulate Cas9 mRNA and sgRNA for gene editing applications. Utilizing IRF4, a transcription factor whose overexpression is linked to myeloma progression and poor prognosis, we co-encapsulated Cas9 mRNA and IRF4 sgRNA. Treatment of myeloma cells with these Cas9 mRNA / IRF4 sgRNA LNPs demonstrated effective IRF4 knockout, which resulted in a cell cycle arrest in the mitotic phase (G2/M), ultimately inducing cancer cell apoptosis. This confirmed the C16-O1 ionizable LNP platform can efficiently deliver functional gene editing cargo, which is particularly important for strategies involving antigen modulation or the knockout of disease-worsening factors.

Intending to translate this LNP for in vivo mRNA delivery to myeloma cells, we utilized maleimide-thiol chemistry to conjugate the C16-O1 LNPs to Fab’ fragments of an anti-human CD38 monoclonal antibody, creating the t-C16-O1 LNP. The successful conjugation was confirmed by dynamic light scattering, which showed a characteristic increase in hydrodynamic diameter of approximately 60 nm. These t-C16-O1 LNPs were administered intravenously in myeloma tumor-bearing mice 21 days after tumor inoculation to evaluate their biodistribution and ability to deliver mRNA LNPs to the tumor site and myeloma cells *in vivo*.

Compared to untargeted C16-O1 and SM-102 LNPs, the t-C16-O1 LNPs demonstrated several key advantages: significantly higher delivery efficacy to the tumor site, a reduced liver-to-tumor delivery ratio (mitigating hepatic clearance), and, crucially, superior functional mRNA delivery specifically to the GFP+ myeloma cells. In conclusion, this study showcases the development of the first targeted LNP system, to our knowledge, for functional mRNA delivery to myeloma cells *in vivo*, establishing a potent new avenue for combination therapies aimed at overcoming current clinical limitations in MM.

