## Supplemental Information for "Anti-CD38-Targeted Piperazine-Derived Lipid Nanoparticles Overcome Hepatic Clearance for mRNA Delivery to Multiple Myeloma Cells *In Vivo*" (from another version of this preprint: version 1, 2025-11-13)

**Contents**

A. **Materials and Methods**
B. **Supplementary Figures**
   1. **Figure S1: LC-MS spectra of lead (C16-O1) ionizable lipid.**
   2. **Figure S2: Structures of C12-200 and SM-102 ionizable lipids.**
   3. **Figure S3: *In vitro* LNP-mediated mCherry mRNA delivery to RPMI-8266, U266, and NCI-H929 CD138+ and CD138- myeloma cells.**
   4. **Figure S4: *In vitro* LNP-mediated Cas9 mRNA / IRF4 sgRNA delivery to RPMI-8266 myeloma cells.**
   5. **Figure S5: *In vivo* mRNA LNP safety via liver enzymatic activity.**
A. **Materials and Methods:**

**Ionizable lipid synthesis.** Ionizable lipids were synthesized by ring-opening conjugation of polyamine headgroups to alkyl epoxide tails, as previously described. Briefly, polyamine cores (Enamine, Kyiv, Ukraine) were reacted with a 10-fold molar excess of alkyl epoxides—1,2-epoxydodecane (C12), 1,2-epoxytetradecane (C14), or 1,2-epoxyhexadecane (C16) (MilliporeSigma, Burlington, MA, USA)—in ethanol under constant stirring at 80°C for 48 h. Following solvent evaporation using a Rotovapor R-300 (Buchi, New Castle, DE, USA), crude products were used directly or further purified. Lipid identity and purity were confirmed by LC-MS (Acquity UPLC C8 column, gradient elution with 1% TFA) and stored as dry films at −20°C until use.

**mRNA production.** Codon-optimized mRNAs encoding mCherry fluorescent protein and Cas9 were synthesized via in vitro transcription (IVT) using T7 RNA polymerase (HiScribe® T7 High Yield RNA Synthesis Kit, New England Biolabs, Ipswich, MA, USA) from linearized plasmid templates. mRNAs were capped with CleanCap AG (TriLink Biotechnologies, San Diego, CA, USA) and contained N1-methylpseudouridine or 5-methoxyuridine modifications, respectively. Transcribed RNAs were purified by LiCl precipitation and resuspended in nuclease-free water. RNA quality was confirmed by agarose gel electrophoresis, and concentration was quantified using a Nanodrop (Infinite 200 PRO, Tecan, Männedorf, Switzerland).

**Lipid nanoparticle (LNP) formulation and characterization.** LNPs were formulated by microfluidic mixing (NanoAssemblr, Precision NanoSystems, Vancouver, Canada) at a 3:1 (aqueous:organic) flow rate. Lipids were combined in ethanol at 35:16:46.5:2.5 molar ratios (ionizable lipid:cholesterol:DOPE:C14PEG2000) and mixed with citrate buffer (10 mM, pH 3.0) containing mRNA at an ionizable lipid-to-RNA ratio of 10:1 (w/w). LNPs were dialyzed in 1x PBS using 10 kDa MWCO cassettes (Thermo Fisher Scientific, Waltham, MA, USA) for 2 h.

Size and polydispersity index (PDI) were measured by dynamic light scattering (DLS; Zetasizer Nano, Malvern Instruments, Worcestershire, UK). For RNA encapsulation, LNPs were diluted 100X in either 1X TE buffer or TE buffer containing 0.1% (v/v) Triton X-100 and incubated for 20 min at room temperature with shaking (350 rpm). Quant-iT RiboGreen reagent (Thermo Fisher Scientific) was added to each well, and fluorescence was measured at 490/520 nm using an Infinite 200 PRO plate reader (Tecan). RNA concentration was calculated using a standard curve, and encapsulation efficiency was calculated as [(Triton – TE)/Triton] × 100.

The apparent pKa was determined using a fluorescence assay with the fluorescent probe 8-anilino-1-naphthalenesulfonic acid (TNS) (MilliporeSigma). LNPs were diluted to a final ionizable lipid concentration of 20 uM in 10 mM citrate buffer, and TNS was added to a final concentration of 1 uM. The solution was titrated across a pH range from 4.0 to 7.0 using either HCl or NaOH. Fluorescence was measured at an excitation wavelength of 320 nm and an emission wavelength of 430 nm (Infinite 200 PRO plate reader, Tecan). The fluorescence data were plotted against the pH and fitted to a sigmoidal dose-response curve to determine the pKa value.

**Targeted LNP synthesis.** For the preparation of targeted LNPs, anti-CD38 antibodies (clone HB-7) were reduced to F(ab')₂ fragments using the Pierce™ Mouse IgG1 Fab and F(ab')₂ Preparation Kit (ThermoFisher Scientific, Cat# 44980), and thiolated with DL-dithiothreitol (DTT, MedChemExpress, Cat# HY-15917). Desalted fragments were added to pre-formed LNPs containing DSPE-PEG-maleimide (Avanti Polar Lipids, Cat# 880126) and incubated at 37°C for 1 hour to allow thiol-maleimide conjugation. Unconjugated antibodies were removed by dialysis in 1X PBS using 1000 kDa MWCO Float-A-Lyzer G2 dialysis devices (Fisher Scientific, Cat# 08-607-030) for 24 hours. Conjugation efficiency was confirmed by DLS.

**Cell culture.** Human multiple myeloma RPMI 8226, U266, and NCI-H929 cells were obtained from ATCC and cultured in RPMI-1640 medium supplemented with 10% FBS and 1% penicillin-streptomycin at 37°C, 5% CO₂. All cell lines tested negative for mycoplasma contamination.

***In vitro* assessment of LNPs.** Cells were seeded in 96-well U-bottom plates and treated with luciferase or Cas9 mRNA-loaded LNPs for 24–72 h. Luminescence was measured using the Bright-Glo Luciferase Assay System (Promega, Madison, WI, USA). For genome editing, Cas9 mRNA LNPs targeting IRF4 were delivered alone or in combination with 1 µM lenalidomide. Cell viability was assessed using CellTiter-Glo (Promega). Cell cycle analysis was performed by propidium iodide staining after ethanol fixation and RNase A treatment, followed by flow cytometry.

**IRF4 gene knockout.** Genomic DNA was isolated 72 h post-treatment using a Monarch® Spin gDNA Extraction Kit (Cat# T3010S, New England Biolabs, Ipswich, MA, USA). The IRF4 locus was PCR-amplified, and editing efficiency was analyzed by Sanger sequencing followed by TIDE analysis.

**Animal studies.** All animal procedures were approved by the University of Pennsylvania Institutional Animal Care and Use Committee (IACUC, Protocol #806540). Female and male CB17/Icr-Prkdcscid/IcrIcoCrl (CB17 SCID, strain 236) mice (6–8 weeks old, Charles River Laboratories, Wilmington, MA, USA) were inoculated subcutaneously with 10 × 10⁶ RPMI 8226 cells in 50% Matrigel (Corning, Cat# 354234). Tumor volumes were measured biweekly with calipers and calculated as (L × W²)/2.

***In vivo* evaluation of LNPs.** Mice bearing palpable tumors were randomized and intravenously injected with targeted or untargeted mRNA LNPs (0.5 mg/kg) encoding mCherry. At indicated timepoints, tumors were harvested for ex vivo imaging or processed to single-cell suspensions for genomic DNA extraction. Luciferase expression was assessed by IVIS imaging (PerkinElmer, Waltham, MA, USA).

For tumor delivery analysis, LNPs were co-formulated with the lipophilic tracer DiR (Thermo Fisher Scientific) at 0.05 mol% of the total lipid content. Tumors were harvested, weighed, and processed into single-cell suspensions using the gentleMACS Dissociator (Miltenyi Biotec). Cell suspensions were analyzed by flow cytometry (CytoFLEX, Beckman Coulter). The total cell populations were gated on viability, and MM cells were identified by their constitutive GFP expression (RPMI-8226 cells). LNP delivery was quantified as the percentage of DiR+ cells within the viable cell gate, and functional delivery was quantified as the percentage of mCherry+ cells within the GFP+ myeloma cell population. All flow cytometry data were analyzed using FlowJo software.

***In vivo* safety assessment.** Serum was collected 6 hours after LNP injection. To assess hepatotoxicity, alanine aminotransferase (ALT; Cayman Chemical, Ann Arbor, MI, USA, Cat# 700260) and aspartate aminotransferase (AST; Cayman Chemical, Cat# 701640) levels were quantified using colorimetric activity assay kits per manufacturer instructions.

**Statistical analysis.** All analyses were performed using GraphPad Prism 10 (San Diego, CA, USA) software; more specifically, statistical analysis was carried out with unpaired two-tailed t-tests or one- or two-way ANOVAs where appropriate. Data were plotted as mean ± s.d. unless otherwise stated.

B. **Supplementary Figures**
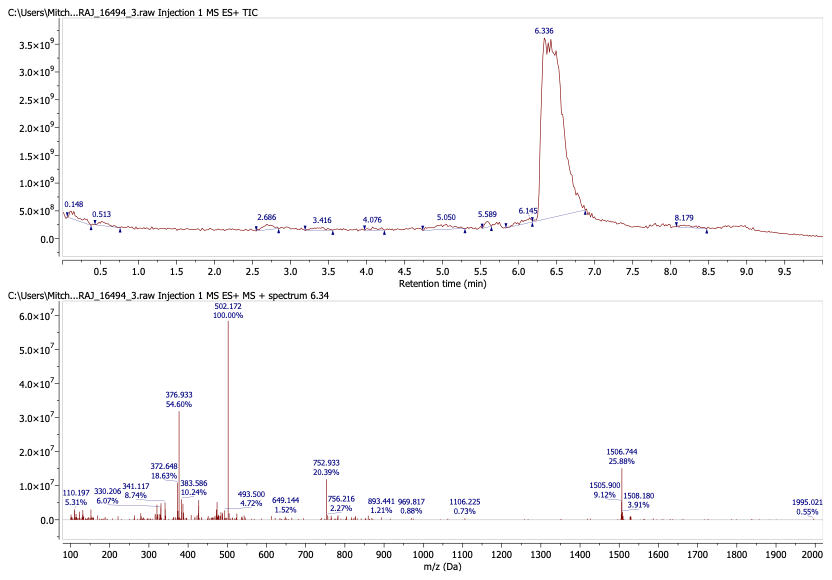

**Figure S1.** LC-MS for the C16-O1 ionizable lipid.

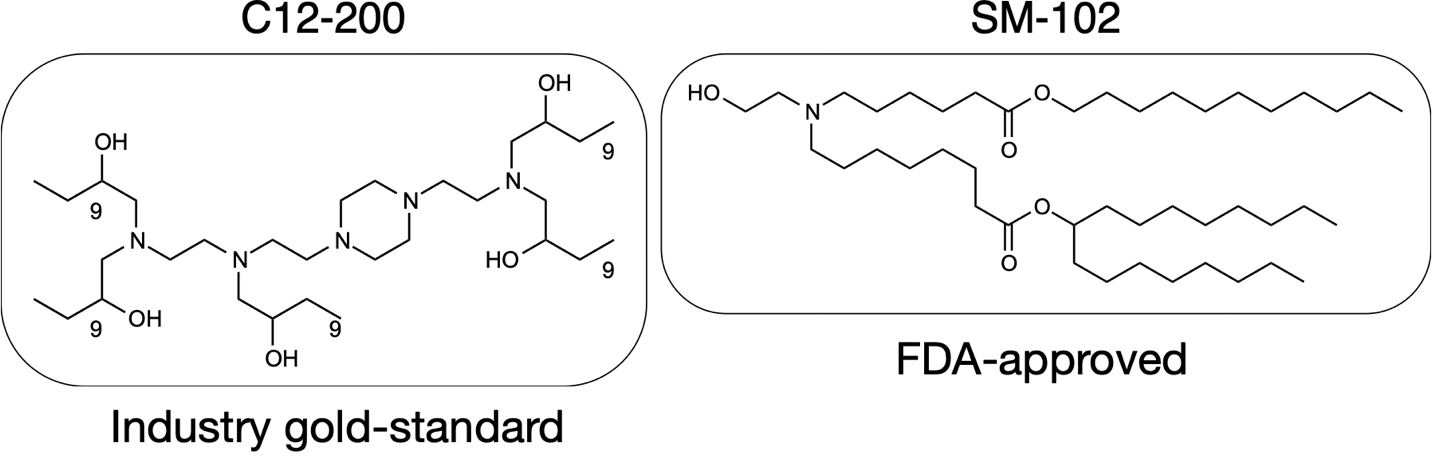

**Figure S2.** Chemical structures of industry standard C12-200 and FDA-approved SM-102 ionizable lipids used in the library as controls.

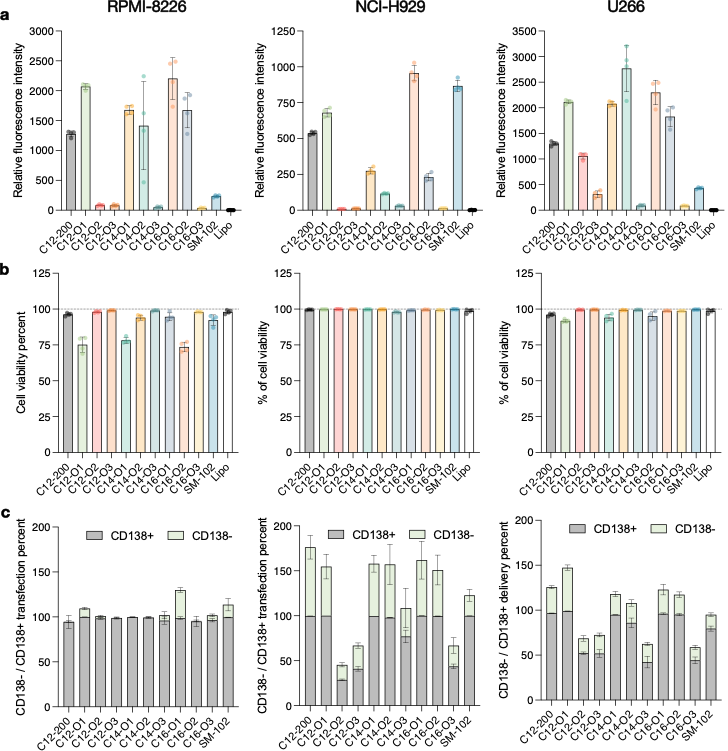

**Figure S3. *In vitro* mRNA-LNP delivery to myeloma cells.** (a) Relative fluorescence intensity obtained when cells were treated with 50 ng of mCherry mRNA per 60,000 cells per well. (b) Cell viability percent to assess safety of LNPs while delivering mCherry mRNA. (c) Percent of CD138+ and CD138- transfected by the different ionizable LNPs. All data plotted are mean ± s.d. All values were normalized to untreated controls. n = 3 biological replicates.

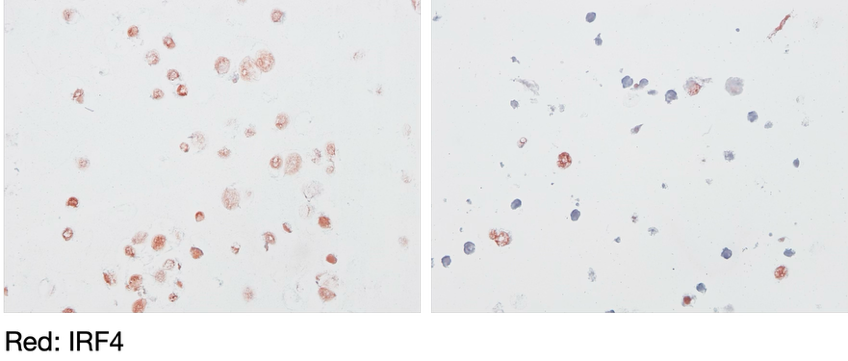

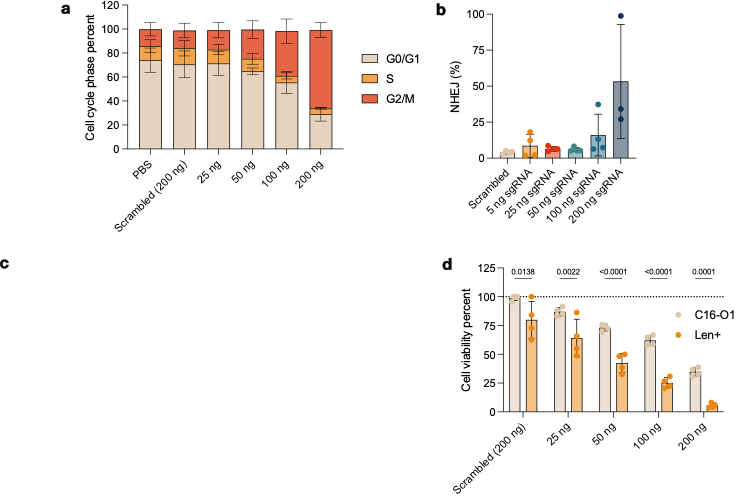
**Figure S4. Cas9 mRNA / sgRNA IRF4 C16-O1 LNPs induce cell-cycle arrest and sensitize myeloma cells to lenalidomide *in vitro*.** (a) Analysis of cell cycle phase in MM cells (RPMI-8226) upon treatment with increasing doses (25 ng to 200 ng of RNA per 60,000 cells) of therapeutic Cas9 mRNA / IRF4 sgRNA C16-O1 LNPs. The data shows a dose-dependent accumulation of cells in the G0/G1 phase, confirming the biological effect of IRF4 knockout. Controls are PBS and scrambled sgRNA (200 ng). (b) Sequencing analysis quantifying the Non-Homologous End Joining (NHEJ) activity in MM cells treated with various doses of IRF4 sgRNA, demonstrating the efficiency of Cas9-mediated gene editing and indel formation. (c) Images of MM cells stained for IRF4 (red chromogen) following treatment with control (left, stron IRF4 signal) versus Cas9 mRNA /IRF4 sgRNA LNPs (right, near-total loss of IRF4 signal). (d) MM cell viability 7 days after treatment with lenalidomide (Len, 10 uM) alone or in combination with increasing doses (25 ng to 200 ng of RNA) of IRF4 sgRNA LNPs. Combination treatment significantly reduces cell viability compared to single-agent treatment, highlighting the therapeutic synergy achieved by genetically disrupting IRF4. Data is plotted as mean ± s.d. n = 3 biological replicates.

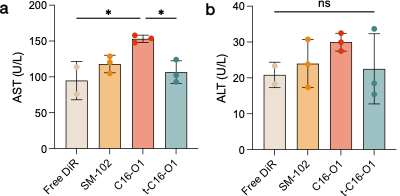

**Figure S5. Assessment of systemic toxicity following intravenous LNP administration.** (a) Quantification of AST in the serum 6 hrs post-injection of the LNP groups (SM-102, C16-O1, and t-C16-O1) compared to the free DiR control. Treatment with the untargeted C16-O1 LNP led to a significant increase in AST levels, indicating transient hepatic stress or injury, which was statistically alleviated in the t-C16-O1 targeted group. (b) Quantification of ALT in the serum 6 hrs post-injection. No statistically significant differences in ALT levels were observed across any of the treatment groups, confirming overall minimal hepatotoxicity associated with the LNPs. Data is plotted as mean ± s.d. n = 3 mice per group.
